# Diamond Raman Laser and Yb Fiber Amplifier for *In Vivo* Multiphoton Fluorescence Microscopy

**DOI:** 10.1101/2021.10.20.464141

**Authors:** Shaun A. Engelmann, Annie Zhou, Ahmed M. Hassan, Michael R. Williamson, Jeremy W. Jarrett, Evan P. Perillo, David J. Spence, Theresa A. Jones, Andrew K. Dunn

## Abstract

Here we introduce a fiber amplifier and a diamond Raman laser that output high powers (6.5 W, 1.3 W) at valuable wavelengths (1060 nm, 1250 nm) for multiphoton excitation of red-shifted fluorophores. These custom excitation sources are both simple to construct and cost-efficient in comparison to similar custom and commercial alternatives. Furthermore, they operate at a repetition rate (80 MHz) that allows fast image acquisition using resonant scanners. We demonstrate our system’s compatibility with fast resonant scanning, the ability to acquire neuronal images, and the capability to image vasculature at deep locations (>1 mm) within the mouse cerebral cortex.

## 1. Introduction

Multiphoton microscopy (MPM) is widely used in neuroscience to visualize neurons and vasculature in mouse models at depths up to approximately 1 mm in the cortical cortex [1-3]. Since MPM relies on the nonlinear absorption of multiple excitation photons to initiate a fluorescent event, ultrafast pulsed lasers must be used [4]. Many strategies to push MPM deeper directly involve modifying the excitation lasers, often addressing their repetition rates or wavelengths.

Titanium-doped sapphire (Ti:S) lasers have long been the standard excitation source for MPM. They typically output femtosecond pulses at an 80 MHz repetition rate with a power and wavelength (700-1000 nm) that can excite many fluorophores efficiently through two-photon absorption. Researchers can effectively image neural tissue to 800 μm depths using these oscillators [5]. To push imaging deeper, many groups have turned to amplifier sources with lower repetition rates and increased pulse energy to improve excitation efficiency without raising average power to levels where thermal damage is risked [6]. Theer et al. used a Ti:S-based regenerative amplifier with a 200 kHz repetition rate to enable imaging to 1 mm in the neocortex [7]. Optical parametric amplifiers (OPAs) are routinely used to image even deeper regions when operating at similar low repetition rates (∼250-500 kHz) [8-10]. This is made possible by their longer wavelength outputs (λ=1100-1900 nm). Photons at longer wavelengths are less susceptible to scattering as they travel through tissue, so the percentage of photons that reach the focal point increases as the excitation wavelength lengthens [11,12]. OPAs are advantageous for deep imaging with their high pulse energies and spectral characteristics, but their low repetition rates can limit imaging speed.

While deep imaging is certainly helpful, so is fast image acquisition. Many new strategies to improve imaging speed have been introduced, including ones that make use of acousto-optic modulators and resonant scanning mirrors [13]. When dramatically increasing the scan speed, it is crucial to ensure the excitation laser’s repetition rate can keep up with the decreased pixel dwell times. Optical parametric oscillators (OPOs) are a long wavelength (often 1100-1600 nm), high repetition rate (80 MHz) option with which Kobat et al. demonstrated imaging well past 1 mm in the mouse brain [14]. These are not without their downsides, however. Custom OPOs are very difficult to build and operate due to their complexity. Commercial oscillators with similar performance to OPOs (Insight, Spectra Physics; Chameleon Discovery, Coherent) can be purchased that have simple turn-key operation, but they come at a significant price. In this paper we demonstrate a cost-efficient, simple alternative involving a fiber amplifier and diamond Raman laser which produce light with high average powers at long wavelengths (1060 and 1250 nm). We demonstrate that the lasers operate at a repetition rate (80 MHz) compatible with resonant scanning strategies. Furthermore, we demonstrate the capability to use our sources for neuronal imaging, and vascular imaging to a depth of 1 mm in the mouse neocortex.

## 2. Materials and methods

### 2.1 Ultrafast Excitation Lasers

The excitation lasers used in this work consist of a fiber amplifier and diamond Raman laser (Fig. 1A and 1B). These are more powerful iterations of similar lasers for which general build instructions are included with a previous publication [15]. For the current fiber amplifier, amplification occurs in the 25 μm core of an ytterbium-doped double-clad polarization maintaining fiber 6 meters in length (Liekki Yb1200-25/250DC-PM, nLight). Custom FC/APC connectors were added to each fiber end to prevent damage when operating at high powers (Coastal Connections, Ventrua, CA). The amplifier is seeded with 75 mW from a commercial fiber oscillator with an 80 MHz repetition rate (Origami, NKT Photonics). The output from a laser diode at λ=915 nm (K915FA3RN-30.00W, BWT) serves as the pump, which is propagated backwards relative to the seed through the 250 μm inner cladding of the fiber. The pump diode power is adjusted to set the output power of the amplifier. Operation in the parabolic amplification regime ensures that pulses acquire only a linear chirp, allowing pulse compression with a transmission grating pair upon output [16-20]. We have achieved an average power as high as 6.5 W following compression, with pulses centered at λ=1060 nm that are approximately 110 fs in length assuming a sech^2^ shape (Fig. 2A-C). Spectra were measured with an NIR spectrometer (AvaSpec-NIR256-1.7, Avantes) and autocorrelations were recorded using a commercial autocorrelator (pulseCheck, APE). The amplifier output is directed to either to the diamond Raman laser or one of our microscopes at a power established using a half-wave plate and Glan-Laser polarizer in combination.

**Fig. 1.**
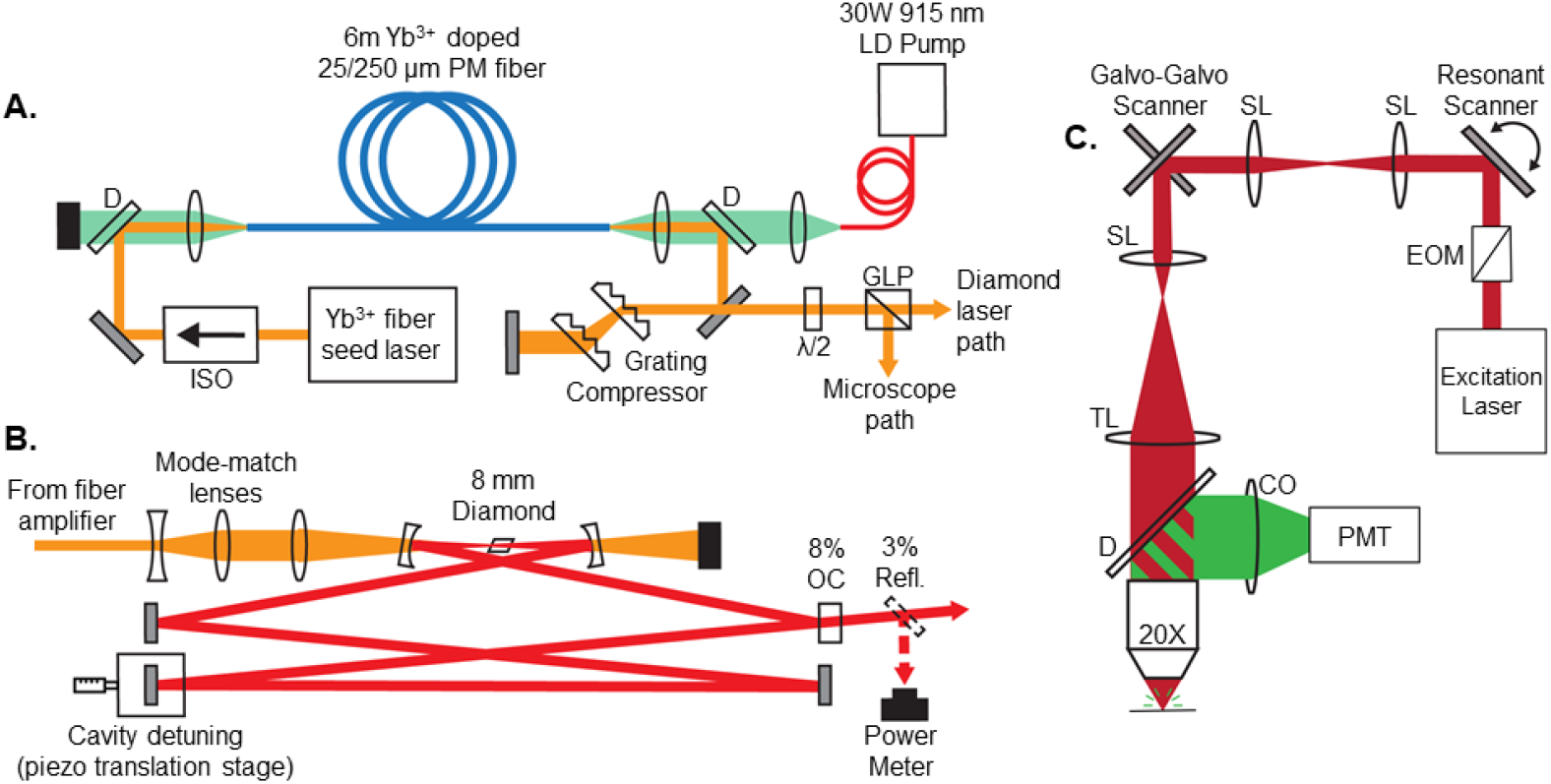
Schematics of the fiber amplifier (A) and diamond Raman laser (B). Also shown in (C) is a schematic of our multiphoton microscope that employed a resonant scanner. D - dichroic, ISO - isolator, λ/2 - half-wave plate, GLP - Glan-Laser polarizer, OC - output coupler, SL - scan lens, TL - tube lens, CO - collection optics, EOM – electro-optic modulator.

**Fig. 2.**
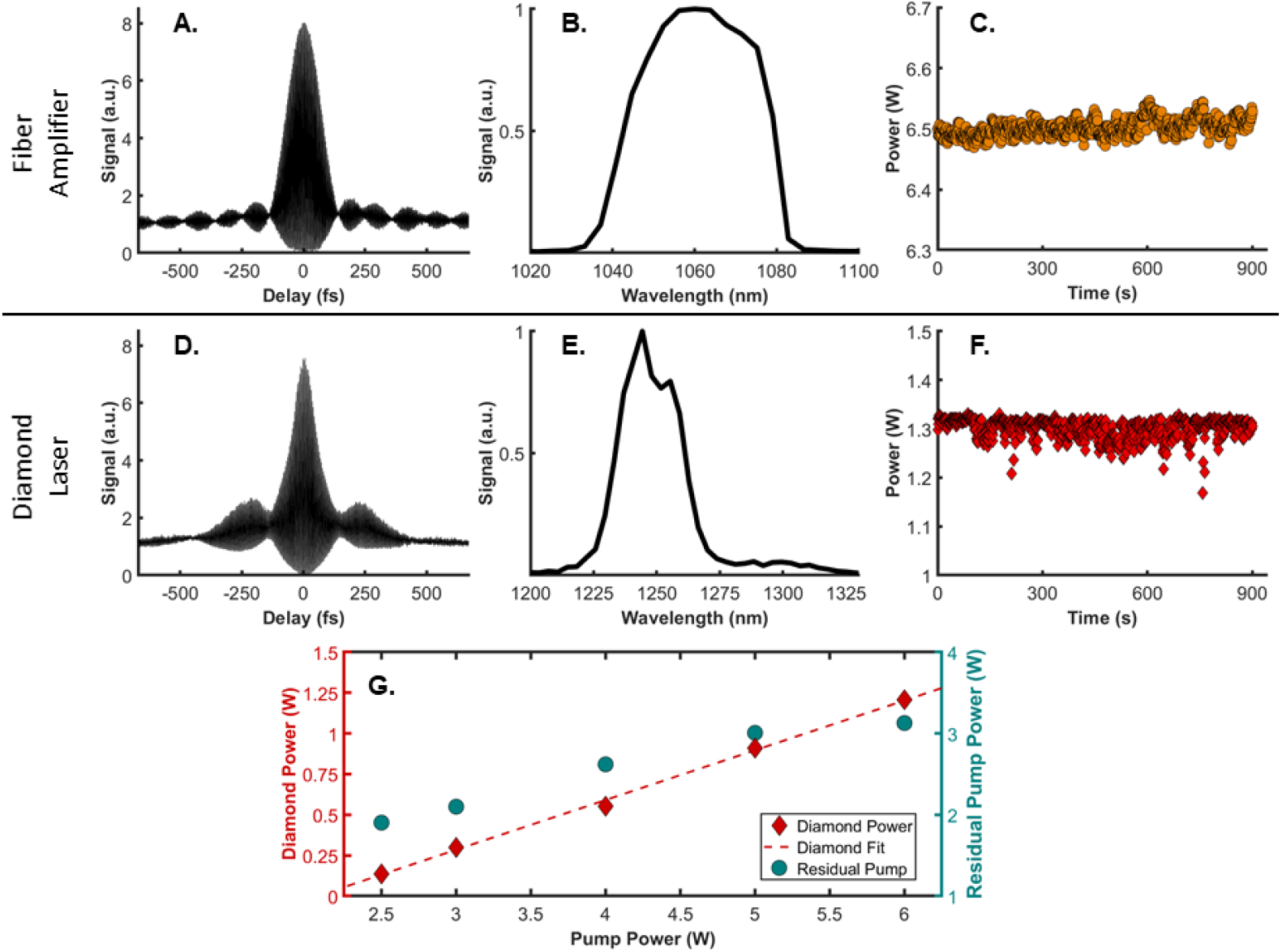
Characterization of laser system. The top row contains the fiber amplifier autocorrelation (A), spectrum (B), and power output over 15 minutes (C). The bottom row contains the same information for the diamond Raman laser (D-F). (F) is the diamond laser output over 15 minutes while pumped with the fiber amplifier at the power shown in (C), using pump pulses possessing a slight negative chirp. (G) displays the diamond laser output power as the pump power from the fiber amplifier is changed. Also included is the power of the residual pump light that exits the diamond laser cavity.

The diamond Raman laser, which is configured as a ring cavity based on previously published designs [21,22], is pumped by the fiber amplifier. Mode-matching lenses expand and then focus the pump to an approximate waist radius of ω_0_=20 μm within the diamond crystal (CVD-Grown, 8mm, AR-coated for 1250 nm). The diamond is positioned using a copper stage that is machined to align the <111> axis of the crystal with the horizontally-polarized pump light from the fiber amplifier. The pump pulses provide Raman gain for a circulating pulse at the Stokes wavelength on each round trip. The ring cavity length matches the pulse separation established by the 80 MHz repetition rate of the fiber amplifier. The cavity is built with high percentage reflectors at λ=1250 nm (R>99.5%) apart from one 8% output coupler that passes Stokes light as the diamond laser output. Cavity length, alignment, and pump pulse width are all balanced to achieve both a stable and high average power output. We found that using pump pulses with a slight negative chirp allows such operation, and that the resulting diamond laser pulses are sufficiently short such that additional dispersion compensation and pulse compression is not needed for our imaging. We have achieved average powers just over 1.3 W with pulses centered at λ=1250 nm that are approximately 100 fs in length with a pump power of 6.5 W (Fig. 2D-F). Fig. 2G shows how the diamond Raman laser output changes with pump power. The resulting slope efficiency is about 31%, comparable with other diamond lasers [21,22]. Also shown in Fig. 2G is the power of residual pump light that exits the ring cavity after passing through the diamond crystal for each pumping condition.

### 2.2 Multiphoton Microscopes

Two multiphoton microscopes were used in this work. In both cases, the excitation laser beam diameter is expanded with a telescope to fill the back aperture of the microscope objective (XLUMPLFLN 20X, 0.95 NA, Olympus). The first microscope will be referred to as the resonant-galvo microscope (Fig. 1C). Here, an electro-optic modulator regulates the beam power sent to the sample (Conoptics, 350-80). A resonant scanner (CRS 8kHz, Cambridge) and a galvanometer scanner (GVS012, Thorlabs) are coupled together with a pair of scan lenses (SL50-2P2, Thorlabs) and are used to steer the excitation light in a raster pattern. An additional scan lens and a Plössl tube lens (AC508-400-C, Thorlabs) image these scanners onto the back aperture of the microscope objective. Fluorescence is epicollected and directed to a photomultiplier tube (H10770PB-40, Hamamatsu) by a 775 nm cutoff dichroic (FF775-Di01, Semrock) and is further filtered by a bandpass filter (FF01-609/181-25, Semrock). Vidrio Technologies’ ScanImage software controlled the image acquisition [23].

The second microscope will be referred to as the galvo-galvo microscope. For this, we now use a half-wave plate in combination with a polarized beam splitter to adjust beam power. A pair of galvanometer scanners (6125HB, Cambridge Technology) steer the excitation light. The previously described scan lens and tube lens combination image the scanners onto the objective back aperture. Once again, a 775 nm cutoff dichroic guides fluorescence to the detector, which is now filtered by a 770 nm shortpass filter (FF01-770/SP, Semrock). The detector here is a PMT with superior sensitivity at longer wavelengths (H10770PB-50, Hamamatsu). Custom software written with National Instrument’s LabVIEW controlled the image acquisition. This microscope was also capable of performing laser speckle contrast imaging [24].

### 2.3 Animal Preparation and Imaging

Animal procedures were approved by The University of Texas at Austin Institutional Animal Care and Use Committee. During both surgeries and imaging sessions mice were anesthetized with isoflurane while body temperature was maintained at 37.5° Celsius. Mice were surgically fit with cranial windows and allowed to recover for at least one week prior to imaging. To label neurons with tdTomato, Ai14 mice (#007914, Jackson Laboratory) were intracortically injected with an AAV5-CaMKII-Cre vector (105558-AAV5, Addgene). Vasculature was labeled through retro-orbital injections of fluorescent dye diluted in physiological saline. Excitation power was minimized when imaging superficial regions and increased with depth. Power incident on the brain surface was maintained under 250 mW to avoid tissue damage [25]. The only instance where excitation powers approached this threshold was during the deep imaging experiments covered in section 3.2.

## 3. Experimental Results

### 3.1 In Vivo Imaging with a Resonant Scanner

Vasculature across a large (>1 mm^2^) lateral field of view was imaged with the resonant-galvo microscope using the fiber amplifier for excitation. The grating compressor was set to minimize the pulse width at the focal plane prior to imaging. 100 μL of 70 kDa dextran-conjugated Texas Red diluted in saline at a 5% w/v ratio was added to the blood plasma prior to the imaging session to serve as the label. 20 frames were averaged at each depth to produce images, which each originally had a 700 by 700 μm field of view. Each stack extended to a depth of 660 μm with a 3 μm step size in between slices. 4 stacks were acquired in adjacent regions and stitched together afterwards using ImageJ [26]. The final stitched mosaic image had a 1160 by 1160 μm lateral field of view. Fig. 3A shows a maximum intensity projection through the entire mosaic stack. Fig. 3B likewise contains a side projection through the stack. A laser speckle contrast image of blood flow on the cortical surface of the imaged mouse is shown in Fig. 3C, where a box was added to indicate the region over which MPM was performed. Matching vascular features are easily observed in both the MPM and the laser speckle contrast images.

**Fig. 3.**
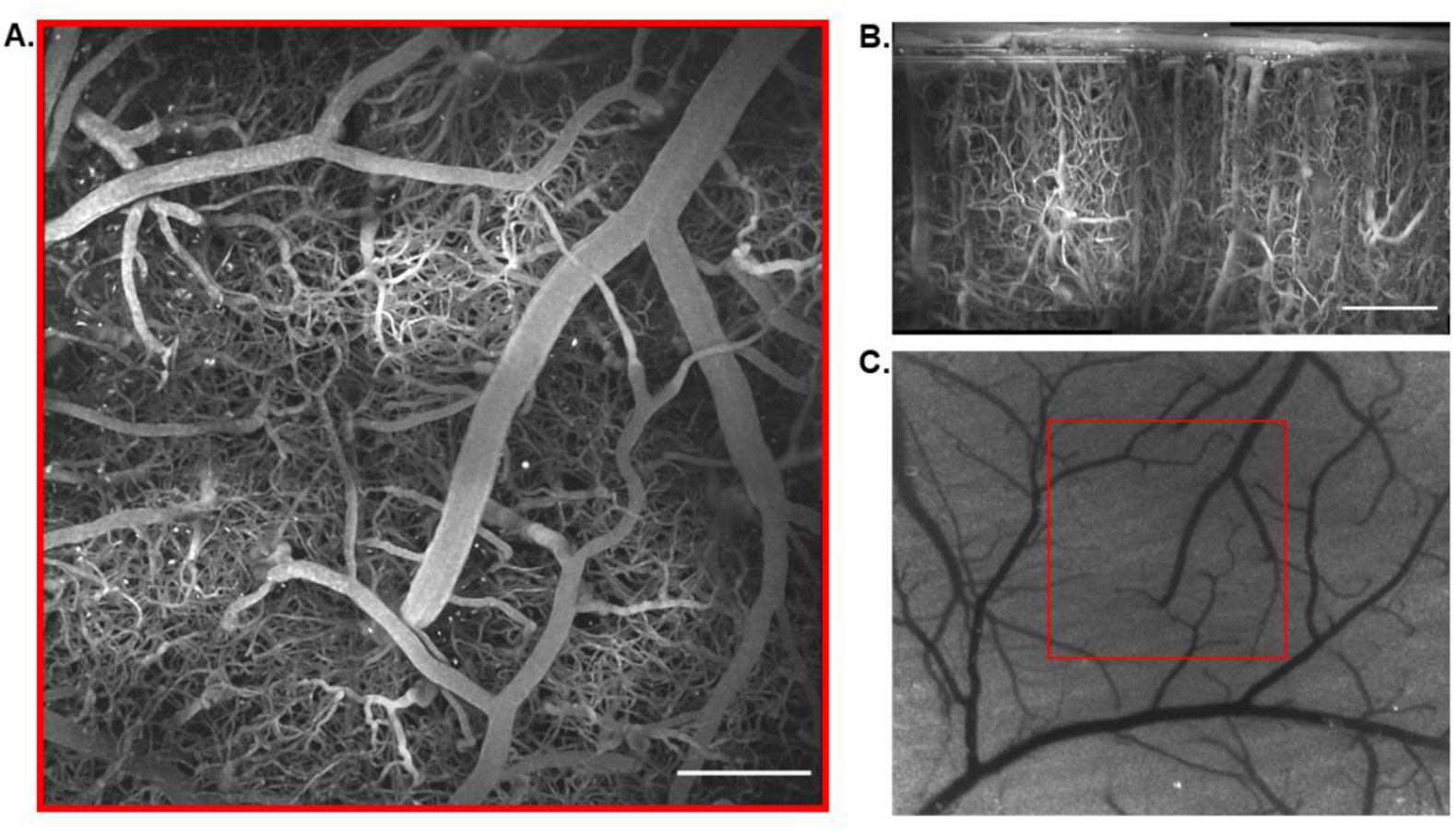
Imaging of Texas Red-labeled vasculature using the fiber amplifier along with a resonant scanner. The lateral field of view is 1160 by 1160 μm. (A) is the overhead maximum intensity projection through the stack, (B) contains the side projection, and (C) is a laser speckle contrast image that depicts the surface vasculature of the imaged brain region. Scale bars are 200 μm.

### 3.2 Deep Imaging with the Diamond Raman Laser

The diamond Raman laser’s spectral properties make it suitable for deep imaging when paired with a red-shifted bright dye. In Fig. 4 vasculature is imaged to just beyond 1 mm in the brain by exciting Alexa Fluor 647 (80 μL injection at a concentration of 5% w/v) and Alexa Fluor 680 (100 μL injection at a concentration of 5% w/v). The Alexa Fluor 647 image stack has a 5 μm step size between slices, whereas the Alexa Fluor 680 stack has a 3 μm step size. Both stacks are composed of images with a 400 by 400 μm field of view. Frame averaging in superficial regions was limited to 3 frames per slice, but this was increased to as high as 10 frames in the deeper regions of the stacks. Excitation power was minimized at the surface and increased with depth, but the average power incident at the brain surface was not allowed to exceed 250 mW. Towards the left of Fig. 4 are the side projections and various overhead maximum intensity projections through the image stacks of vasculature labeled with Alexa Fluor 680. Towards the right of Fig. 4 is the same content for the vasculature labeled with Alexa Fluor 647. Note the large penetrating vessel that appears at a depth of about 500 μm for the Alexa Fluor 680 images. If this did not appear, we would expect to resolve the vasculature towards the left side of the projection from 900 to 1056 μm just as well as on the right.

**Fig. 4.**
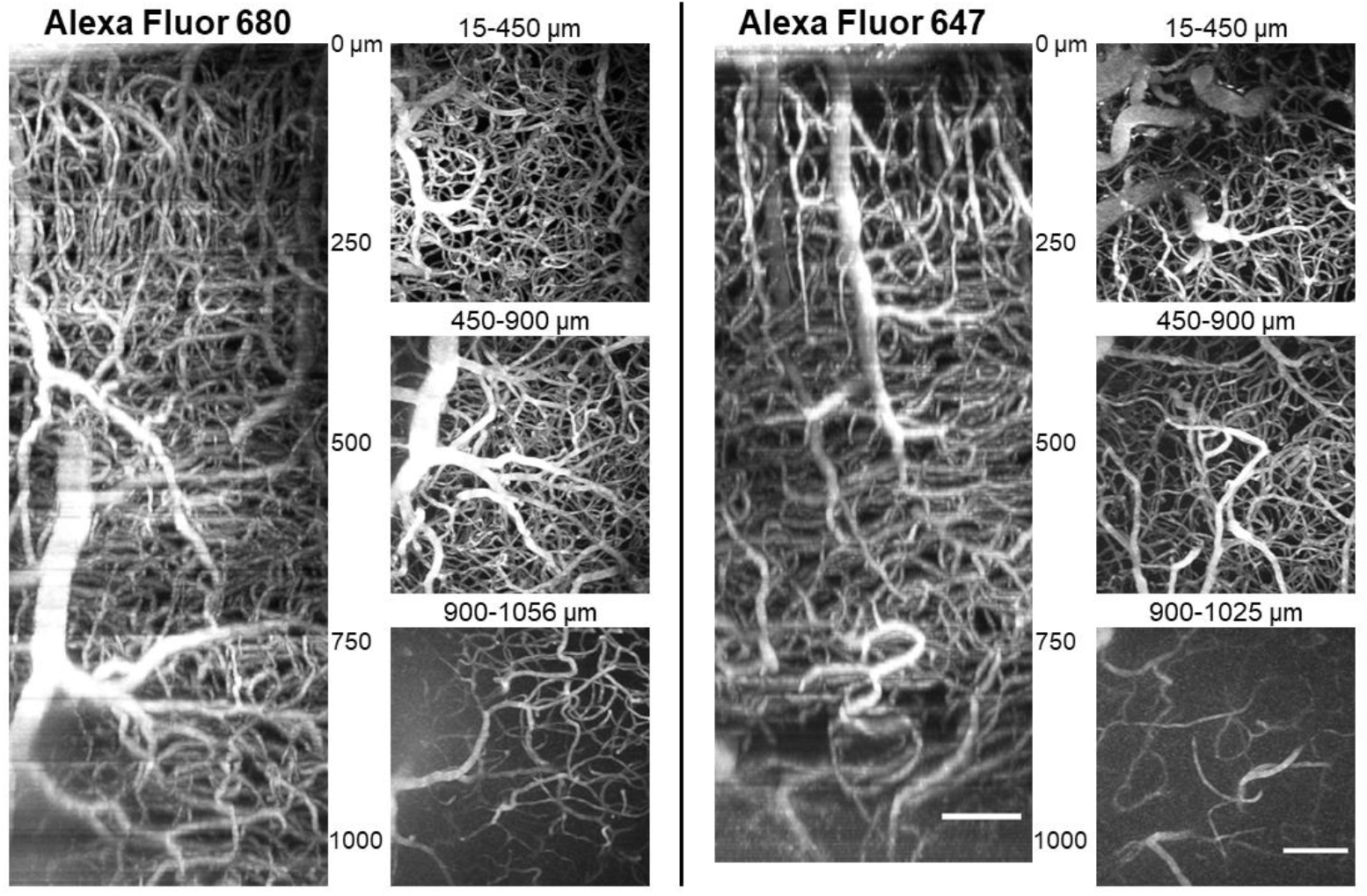
Deep vascular imaging of Alexa Fluor 680 (left) and Alexa Fluor 647 (right) with the diamond Raman laser. Both side projections and overhead maximum intensity projections are displayed. The lateral field of view for the images is 400 by 400 μm. Scale bars are 100 μm.

### 3.3 Neuronal and Vascular Imaging

Neuronal and vascular images from the same cortex location in a mouse were acquired separately during a single session using the galvo-galvo microscope (Fig. 5). Pyramidal neurons expressing tdTomato and vasculature labeled with an 80 μL retro-orbital injection of 10 kDa dextran-conjugated Alexa Fluor 647 diluted in saline at a 2.5% w/v ratio were imaged. Signal from one fluorophore versus the other could be discerned based on the excitation source since Alexa Fluor 647 is efficiently excited by only the diamond Raman laser at λ=1250 nm and tdTomato is efficiently excited by only the fiber amplifier at λ=1060 nm. First, vascular images were acquired using the diamond Raman laser. Images were recorded to a depth of 750 μm using a 3 μm step size between slices, and 3 frames were averaged to create each image in the stack. Imaging of the tdTomato-labeled neurons using the fiber amplifier as the excitation source followed the acquisition of the vascular stack. Images were acquired to a depth of 750 μm with a 3 μm step size, and 5 frames were averaged together at each depth. The lateral field of view for the vascular imaging was ∼400 by 400 μm, whereas the neuronal imaging field of view was 235 by 235 μm. The vascular images shown in Fig. 5 have been cropped to match the neuronal field of view. The two stacks were merged using MATLAB. Fig. 5A shows maximum intensity projections from the surface to a depth of 450 μm for the neuronal stack, the vascular stack, and the merged stack. Fig. 5B contains side projections through the entire stacks for each structure. Axons stemming from the neuronal cell bodies are clearly seen in the side projections. Fig. 5C contains some projections through various depths of the merged stack. Dendrites can be seen stemming from neurons towards superficial regions, and neuronal cell bodies can be seen throughout the entire depth.

**Fig. 5.**
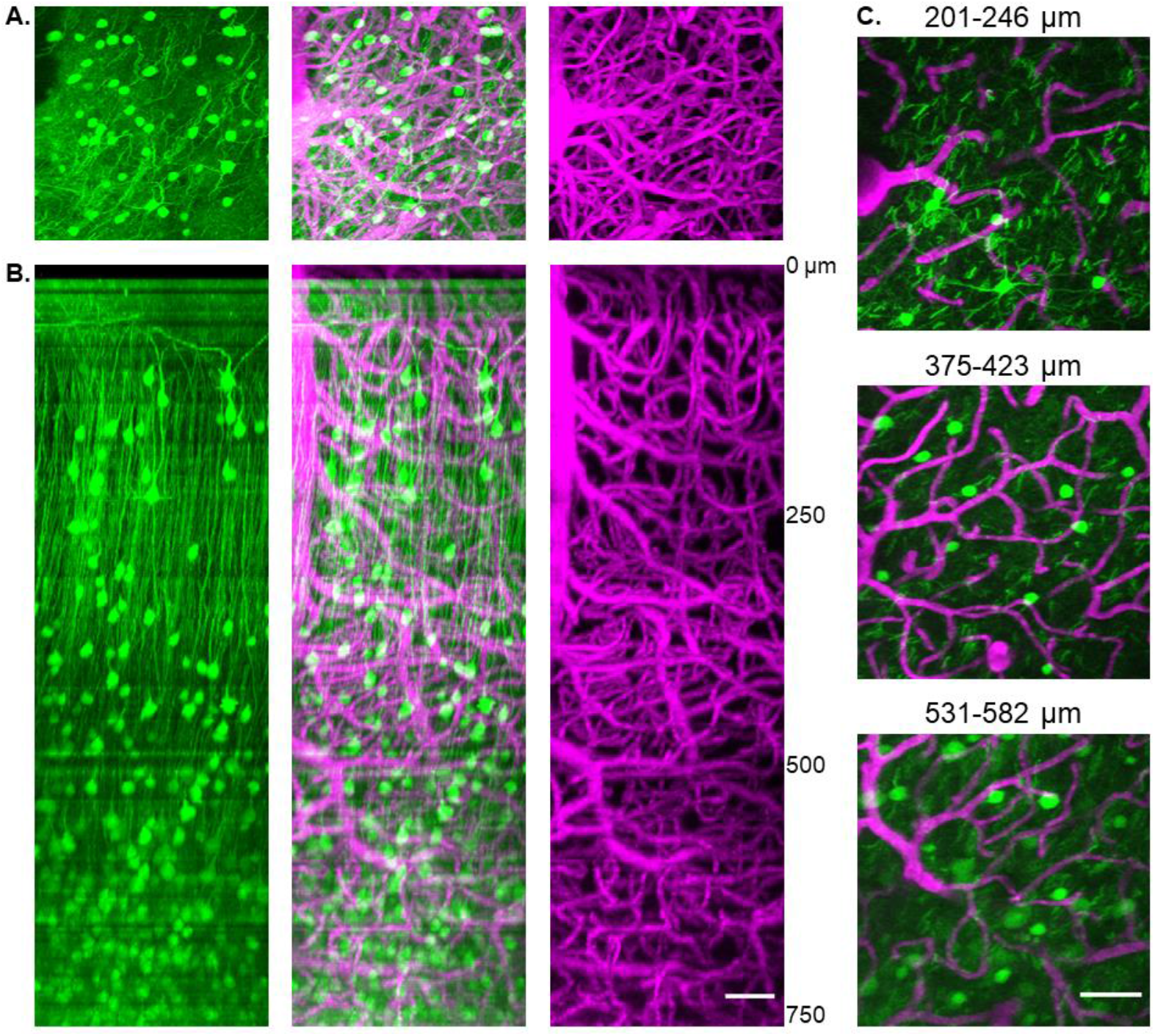
Neuronal and vascular images. (A) Maximum intensity projections spanning from the surface to a depth of 450 μm for neurons (left, 1060 nm excitation), vasculature (right, 1250 nm excitation) and the combination (middle). (B) side projections for the same stacks from (A). (C) Maximum intensity projections at different depths within the stack. Scale bars are 50 μm.

## 4. Discussion

The fiber amplifier and diamond Raman laser presented here offer many unique opportunities. Most notably, they produce high average powers with a relatively simple design that is straightforward to implement. The overall design is similar to our previous work [15] and the modifications made did not significantly add to the system cost. It is still ∼$50,000 for all components, including the seed laser. The new design has increased output power for the fiber amplifier and the diamond laser respectively to 6.5 W at λ=1060 nm and 1.3 W at λ=1250 nm. These power and wavelength combinations allow deep imaging while operating at a repetition rate (80 MHz) that enables fast image acquisition using resonant scanners. When adopting a fast scanning strategy it is important to ensure the excitation laser has a repetition rate that delivers multiple pulses within the shortened pixel dwell time. In Fig. 3 we demonstrate that our lasers can do this when paired with our 8 kHz resonant scanner. Note that 20 frames were averaged together in the images acquired for this figure, while we limited averaging to 3 frames for similar images acquired using galvanometer-driven scanning. Despite this it is still more than 3 times faster to record stacks with the resonant scanner. Decreasing imaging time reduces the possibility of damaging the brain through prolonged exposure to excitation light or harming mice through excessive anesthetic inhalation. It also reduces the magnitude of signal reduction throughout a vascular imaging session due to fluorophore clearance from the bloodstream. Addressing these issues may prove beneficial for chronic studies involving a large field of view that are aided by resonant scanning. Note that while we did not test the diamond Raman laser with resonant scanning, we do not foresee any issues with this.

While we did not use the diamond Raman laser with resonant scanning, we did make use of this laser to image relatively deep brain regions. Both excitation sources in our system produce pulses compressible to relatively short pulse widths (∼100 fs). This, along with their high average powers (both >1W) and relatively long wavelengths, enables deep imaging which we demonstrate in Fig. 4. The diamond Raman laser was used to image vasculature to a depth of 1 mm. Note that we did not use a compressor to optimize the pulse width at the imaging plane for this, which could enable even deeper imaging. We decided that it was not necessary to add dispersion compensation upon determining we could image through the entire depth of the cerebral cortex (∼1 mm) without it.

When compared to other common high repetition rate excitation sources such as Ti:S lasers, both the fiber amplifier and diamond Raman laser have longer wavelength outputs. To take advantage of the decreased scattering offered by the long wavelengths of our lasers, fluorophores that are efficiently excited by each source must be identified. The λ=1250 nm output of the diamond Raman laser pairs well with both Alexa Fluor 647 and Alexa Fluor 680 as shown in Fig. 4. The λ=1060 nm output of the fiber amplifier pairs well with Texas Red for vascular imaging (Fig. 3), and tdTomato which we used to label neurons (Fig. 5). Also important, the spectral difference between the two lasers is sufficient such that the fluorophores excited by one source are not easily excited by the other. This allows us to image multiple structures in a single sample if they are labeled with different fluorophores. As an example, we separately imaged vasculature labeled with Alexa Fluor 647 and then neurons labeled with tdTomato during the same session to create the merged images shown in Fig. 5. In addition to the labels previously mentioned, there is currently a push to develop red-shifted fluorophores that are excitable by far-red wavelengths like what our lasers provide [27-29]. The utility of the laser system will expand as new labels are introduced. Our system can also excite fluorophores at intermediate effective wavelengths if used to initiate non-degenerate excitation (NDE) [15,30-32].

One limitation of the laser pair presented here is their fixed wavelengths. Alternatives such as OPOs and commercial laser systems (Insight, Spectra Physics; Chameleon Discovery, Coherent) have tunable outputs (∼1100-1600 nm for OPOs, 700-1300 nm for commercial systems) and similarly high repetition rates. In practice shorter wavelengths are less useful for deeper *in vivo* imaging due to increased scattering, and wavelengths between 1400 and 1600 nm are rarely used for MPM due to increased water absorption in tissue [12]. Tuning an OPO to a specific wavelength can maximize excitation efficiency for a given fluorophore, but this ability adds complexity to the instrumentation. When using a custom OPO, exact phase matching conditions must be set to produce a desired wavelength. The fiber amplifier and diamond laser system is much simpler in that it is for the most part turn-key. Building the laser system is not significantly more complex than building a custom multiphoton scanning microscope. After initial alignment only the diamond laser’s cavity length needs to be routinely adjusted. Therefore, the system presented in this paper is a cost-efficient, long wavelength multiphoton excitation source with high output powers that can be constructed by labs with some optics expertise.

## 5. Conclusion

MPM is rapidly improving as new technologies and laser systems for the imaging strategy are developed. We have introduced a set of custom excitation lasers for MPM that are valuable for imaging neural structures. The lasers are a fiber amplifier and a diamond Raman laser with high power outputs (6.5 W, 1.3 W) at wavelengths (1060 nm, 1250 nm) excellent for exciting many red-shifted neural labels. We have shown that neurons and vasculature in the mouse cerebral cortex can be reliably imaged with the system. Vascular structure was imaged to depths greater than 1 mm. Additionally, enabled by their high repetition rate (80 MHz), we have shown that our lasers are compatible with a resonant scanning technique that allows fast image acquisition. The laser system we developed is a simple and cost-efficient alternative to many other common custom and commercial excitation sources.

